# Associations of human femoral condyle cartilage structure and composition with viscoelastic and constituent-specific material properties at different stages of osteoarthritis

**DOI:** 10.1101/2022.05.01.490026

**Authors:** Mohammadhossein Ebrahimi, Aleksandra Turkiewicz, Mikko A.J. Finnilä, Simo Saarakkala, Martin Englund, Rami K. Korhonen, Petri Tanska

## Abstract

The relationships between structure and function in human knee femoral cartilage are not well-known at different stages of osteoarthritis. Thus, we characterized the depth-dependent composition and structure of normal and osteoarthritic human femoral condyle cartilage (*n* = 47) and related them to their viscoelastic and constituent-specific mechanical properties. We observed that, in superficial cartilage, the collagen network disorganization and proteoglycan loss were associated with the smaller initial fibril network modulus (collagen pretension). Furthermore, the proteoglycan loss was associated with the greater strain-dependent fibril network modulus (a measure of nonlinear mechanical behavior). The proteoglycan loss was also associated with greater cartilage viscosity at a low loading frequency (0.005 Hz), while the disorganization of the collagen network was associated with greater cartilage viscosity at a high loading frequency (1 Hz). Our results suggest that proteoglycan degradation and collagen disorganization reduce the pretension of the collagen network while proteoglycan degradation also increases the nonlinear mechanical response of the collagen network. Further, the results also highlight that proteoglycan degradation and collagen disorganization increase the viscosity of cartilage, but their contribution to increased viscosity occurs in completely different loading frequencies.

## 1. Introduction

Human articular cartilage plays a vital role in providing a smooth and frictionless articulating surface inside the joints. Its multiphasic nature dictates the tissue’s nonlinear time-dependent mechanical response. The tissue is made up mainly of interstitial fluid, whereas its solid phase consists of proteoglycans (PGs) and collagen fibrils. The constituents of the tissue and their interactions determine the mechanical behavior of the tissue. The negatively charged PG matrix withstands compressive loads when tissue undergoes prolonged loading, while both the pressurized interstitial fluid and collagen network mainly resist instantaneous and dynamic loadings ^14,15^.

Osteoarthritis (OA) is a common joint disease that includes the degradation of articular cartilage, and it affects the mechanical function of the tissue and the joint itself. The disease diminishes hyaline cartilage integrity by affecting its macromolecules. For instance, disorganization of collagen fibers, loss of PGs and collagens are among the few well-known consequences of the disease ^11,27,47^. These alterations weaken the function of the tissue and contribute to the disease progression.

To understand the OA development, several studies with animal and human cartilage have attempted to relate the composition and structure of the tissue to its mechanical properties ^45,49^. Animal models are accessible candidates used to establish the structure-function relationships of normal and osteoarthritic cartilage. The disease can be conveniently induced in animals, where factors such as body mass index, age and daily activity can be controlled ^8,22,48^. Studies performed on human tissue, however, are limited due to accessibility, ethical issues and pricey harvesting process, yet the differences between human and animal tissues have been acknowledged in the literature ^3,7,35^. For clinical translation, it is, therefore, crucial to understand how the properties of human cartilage are altered during the disease progression.

Regarding studies performed on human tissue, the structure-function relationships of healthy and diseased tissue have been studied only in tibial ^15^, patellar ^38^ and hip joint ^34^ cartilage. Nevertheless, structure-function relationships of human femoral condyle cartilage are still not well explored. It is worth noting that the composition and structure of cartilage might differ between joint sites due to their adaptation to different loading environment ^18,53^. Moreover, due to site-specific variations in tissue composition and the loading regime, the disease progression might be different. Hence, it is necessary to investigate human articular cartilage in different sites.

Computational finite element modeling coupled with sophisticated material models developed to capture the complex nonlinear and time-dependent behavior of cartilage offers a possibility to separate the contribution of tissue constituents on the mechanical function of the tissue. Among these material models, the fibril-reinforced poroelastic (FRPE) model has demonstrated a great match with experimental data obtained from cartilage ^14,29,31^. In that model, the mechanical behavior originating from the non-fibrillar matrix (e.g., PGs), fibrillar matrix (e.g., collagen network) and fluid flow are explicitly accounted for. The fluid pressurization and the fibrillar network modeled in the FRPE material model can accurately resemble the peak forces observed in experiments during rapid loading, while the nonfibrillar matrix primarily accounts for the cartilage stiffness in the equilibrium phase. By relating the constituent-specific material parameters to the tissue composition and structure in different stages of OA, one can determine which constituent impairs the mechanical function of the tissue, for instance, during the disease progression. This may also help in linking changes in the mechanical function of the tissue to tissue mechanobiology.

In our recent study ^13^, we investigated human femoral condyle cartilage and characterized the alterations in the bulk mechanical as well as constituent-specific mechanical properties of the tissue with regard to different stages of OA. Surprisingly, we observed no essential deterioration in the mechanical properties of the moderate OA cartilage as compared with that of normal cartilage, while the severe OA cartilage showed clear signs of deterioration. The former observation contradicted the findings obtained from human tibial cartilage ^14^. Hence, in the present study, we aimed to explore the relationships between structure, composition and function to explain our observations on the constituent-specific FRPE and viscoelastic mechanical properties of femoral condyle cartilage at different stages of OA. We hypothesize that the lack of deterioration in the mechanical properties of normal and moderate OA cartilage is because our moderate OA samples have maintained the condition of their collagenous and PG matrices present in normal cartilage. This hypothesis was tested through microscopic and spectroscopic characterization of cartilage structure and composition and associating sturture and composition to the constituent-specific and viscoelastic mechanical properties.

## 2. Materials and Methods

### 2.1. Sample preparation, mechanical testing and determination of the elastic, viscoelastic and constituent specific material parameters

We retrieved 47 osteochondral samples from medial and lateral femoral condyle of 15 total knee replacement (TKR) patients (age 65.0 ± 7.3 years, range 50–79 years, sex: 7 males and 8 females, Trelleborg Hospital, Trelleborg, Sweden) with end-stage medial compartment knee OA and 10 healthy donors (age 53.4 ± 17.8 years, range 18–77 years, sex: 5 males and 5 females, Skåne University Hospital, Lund, Sweden). The general workflow of the study is presented in Figure 1. The biomechanical properties of the samples were characterized in our previous study^13^. We were not able to measure 12 samples in the indentation test due to the lack or considerable loss of articular cartilage (nearly complete erosion). As a result, we included only 35 samples in the mechanical characterization. Since the mechanical testing protocol and results were reported in detail in our recent papers ^13,14^, only a brief summary is given here.

**Figure 1:**
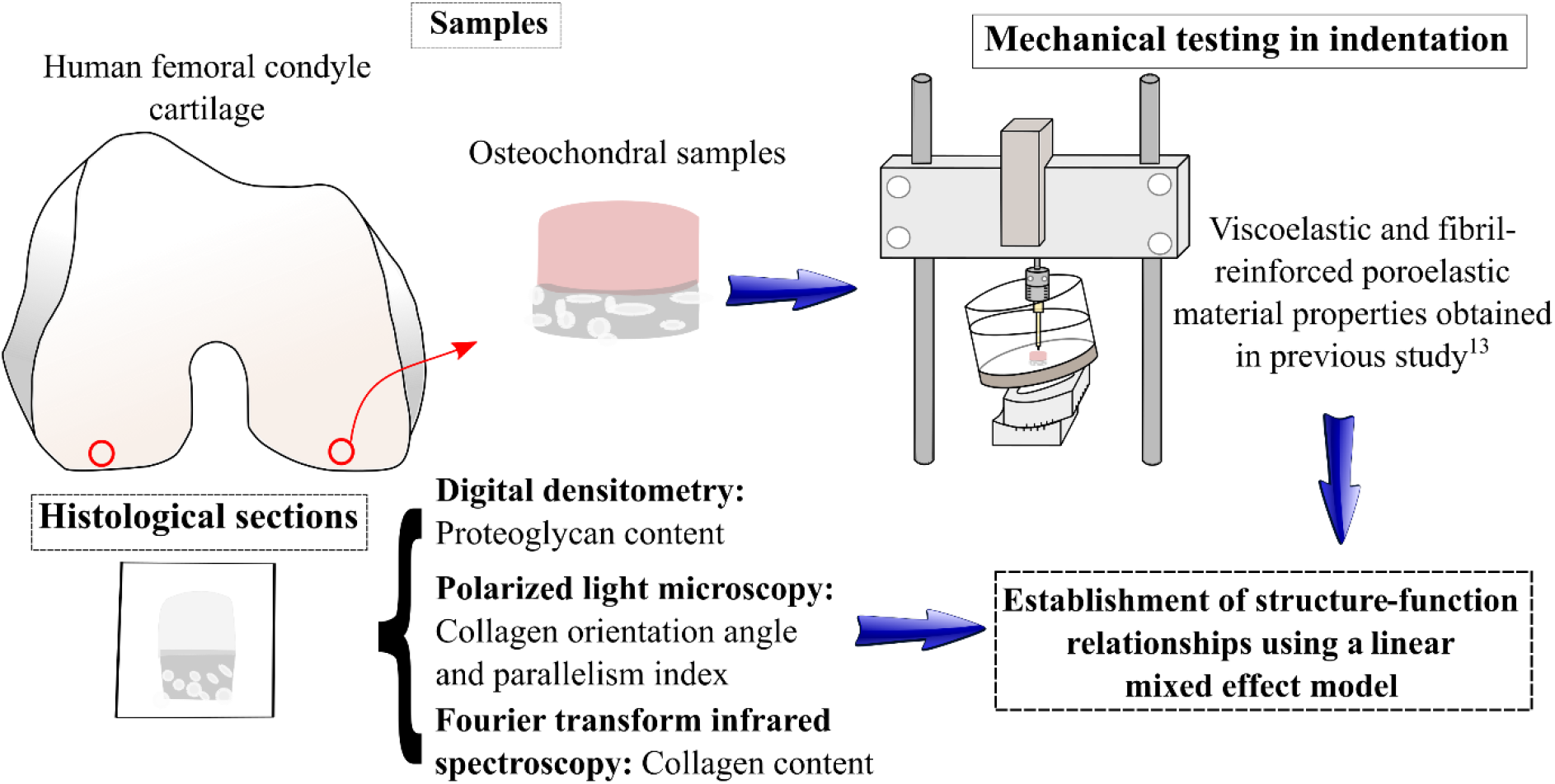
Workflow of the study. First, biomechanical measurements were conducted for the osteochondral samples obtained from tibia. Then, the samples were formalin-fixed, decalcified and embedded in paraffin tissue blocks. The blocks were sectioned and stained for OARSI grading. These steps were conducted in our previous study 13. In this study, the blocks underwent additional quantitative histological analyses including digital densitometry, polarized light microscopy and Fourier transform infrared spectroscopy.

Mechanical indentation testing using a 4-step stress-relaxation protocol followed by a dynamic test with several frequencies was performed (indenter diameter = 0.73 mm). The viscous parameter (phase difference between force and displacement) was calculated accordingly. Afterwards, the finite element models of the samples were constructed and cartilage mechanical behavior was simulated using the FRPE material model. The unknown FRPE material parameters (i.e., the initial fibril network modulus, strain-dependent fibril network modulus, non-fibrillar matrix modulus, initial permeability and strain-dependent permeability coefficient) were obtained by fitting the finite element model’s stress-relaxation response to the corresponding experimental response. After the mechanical testing, the samples were graded using histopathological OARSI grades and grouped to normal (OARSI 0-1.5), moderate OA (OARSI 2-3.5) and severe OA (OARSI 4-5.5) groups. In the current study, the sample set of the previous study (*n* = 35) and the additional sample set, that was unmeasurable in the mechanical indentation test (*n* = 12), were processed to determine quantitative compositional and structural properties. This study was approved by the regional ethical review board of Lund University (Dnr 2015/39 and Dnr 2016/865).

### 2.2. Depth-dependent microscopical and spectroscopical analyses

The structure and composition of cartilage samples were characterized throughout their depth. Following the mechanical testing, the samples were fixed in formalin, decalcified, dehydrated in graded ethanol solutions and embedded in paraffin. Thin histological sections were prepared by microtome from the middle of the sample and perpendicular to the surface. This area matches the area in which the biomechanical indentation test was performed. The slice thicknesses were as follows: 3 µm for digital densitometry (DD), and 5 µm for both polarized light microscopy (PLM) and Fourier transform infrared spectroscopy (FTIR). Three sections were prepared per each sample. Paraffin was removed before measurements.

Safranin-O, known to stoichiometrically bind to negatively charged PGs ^46^, was used to stain sections to be measured by DD. Stained sections were imaged using a conventional light microscope (Nikon Microphot FXA, Nikon Co., Tokyo, Japan. pixel size = 3.09 × 3.09 µm), a CCD camera (Hamamatsu ORCA-ER, Hamamatsu Photonics, Hamamatsu, Japan) and a monochromator (λ = 492 ± 5 nm) ^10,40^. The device was calibrated using neutral filters with a wide range of optical densities (0 to 3). The optical densities acting as an indicator of the stain intake were measured for each sample and used as an estimation of the PG content.

To determine the collagen fibril architecture, a PLM device was used to estimate the main orientation of the collagen fibrils in each pixel. We used a conventional light microscope (Nikon Diaphot TMD, Nikon, Inc., Shinagawa, Tokyo, Japan, pixel size = 2.53 × 2.53 µm) equipped with Abrio PLM system (CRi, Inc., Woburn, MA, USA) to determine the collagen orientation angle. The orientation angle was determined based on Stokes parameters as thoroughly explained in Rieppo et. al. ^44^. As Abrio PLM system does not provide raw data to calculate the tissue parallelism index (i.e., indicating the tissue anisotropy), we employed a custom-made PLM device to calculate the parallelism index, also known as anisotropy index. The custom-made PLM device included a light microscope (Leitz Ortholux II POL, Leitz Wetzlar, Wetzlar, Germany) coupled with a monochrome camera (BFS-U3-88S6M-C FLIR Blackfly® S, FLIR Systems Inc., Wilsonville, OR, USA, pixel size 3.5 × 3.5 µm) equipped with a monochromatic light source (Edmund Optics Inc., Barring-ton, NJ, USA, λ = 630 ± 30 nm), two crossed polarizers (Techspec optics® XP42-200, Edmund Optics, Barrington, NJ, USA) and a stepper motor rotator mount (rotation step = 9°, Newport PR50, Irvine, CA, USA,). The parallelism index (*PI)* was calculated using Michelson’s contrast method ^44^. For a sample with perfectly parallel fibrils (anisotropic structure) *PI* = 1 and fully random fibrils (isotropic structure) *PI* = 0. The collagen orientation angle and fibril parallelism index were obtained for each sample to indicate the collagen architecture.

FTIR device uses infrared light to determine the absorption spectrum representing the molecular content of the tissue. We employed Agilent Cary 600 spectrometer coupled with Cary 610 FTIR microscope (Agilent Technologies, Santa Clara, CA, USA, pixel size = 5.5 × 5.5 µm, spectral resolution = 4 cm^-1^) to measure an infrared spectrum of cartilage samples from the wavenumber range 3800-750 cm^-1^. Then, we applied a constant baseline correction for the spectrum ranging from 2000 to 900 cm^-1^ and, subsequently, calculated the area under the Amide I peak (1720-1595 cm^-1^) known to represent collagen content ^5,32^.

The measured compositional and structural data (i.e., PG content, collagen content, collagen orientation angle and tissue anisotropy) were analyzed using a rectangular region of interest (∼1.5 mm wide throughout the whole tissue thickness) to obtain depth-wise profiles averaged over 3 sections (Matlab software v7.10.0, Mathworks Inc., MA, USA).

### 2.3. Statistical analyses

To estimate the differences in the depth-dependent value of proteoglycan content, collagen content and fibril parallelism index between the three OA groups, we used a linear mixed effects model. To handle the nonlinear shapes of the parameters over depth, we included cubic splines (with knots at 5, 15, 30, 50, 85 and 95 angle degrees), the OA group and their interaction (to allow for depth-specific between-group differences), and compartment as fixed effects. The compartment and individual were included as random effects, compartment nested within an individual. To analyze the differences in the median of collagen orientation (variable bounded by values of 0 and 90 degrees) we used quantile regression for clustered data, with the categorized depth (10 strata for each 10% of thickness), OA-group, their interaction and compartment as fixed effects. We used cluster standard errors to account for the dependence of the data coming from the same individual. We used the quantile regression and chose to compare medians due to the non-normal, heteroscedastic and bounded nature of the collagen orientation variable. We used similar models to estimate differences between the groups in the superficial zone only. The outcome variables were averaged over the superficial 10% of the thickness and thus no splines were used. The proteoglycan content was log-transformed to stabilize the variance.

To analyze the association between the structural variables, constituent-specific mechanical parameters and phase differences, we used a linear mixed regression model. To obtain adequate model fit (i.e., where the residuals are normally distributed, homoscedastic and the relationship between predictors and outcomes is linear), we log-transformed the outcomes: the non-fibrillar matrix modulus, strain-dependent fibril network modulus and initial permeability. Further, we used logit transformation for the collagen orientation angle and log transformation for proteoglycan content. We standardized all predictors (i.e., structural/compositional variables) so that the estimated coefficients represent the difference in the outcome per 1 standard deviation (SD) of the predictor. All four structural/compositional variables and compartment were included in the model as independent variables. An individual was included as a random effect. These models were fitted using structural variables averaged over the bulk cartilage (0-100% of thickness) or the superficial cartilage (0-10% of thickness). We reported relevant estimates with 95% confidence intervals (CIs). The analyses were performed using Stata (v17, College Station, TX, USA).

## 3. Results

The descriptive data over the study sample (age, sex and BMI), including the OARSI scores per donor/TKR and the compartments are presented in Supplementary Table S1 for the analyses of depth-wise composition/structure comparisons between OA groups (*n* = 47) and in Supplementary Table S2 for the analyses of structure-function relationships (*n* = 35).

### 3.1. Depth-wise structural and compositional alterations of human femoral condyle cartilage at different stages of OA

We observed a minor loss of superficial PG content when the moderate OA group was compared to the normal group (4-30% of tissue thickness, Figure 2A). The severe OA group showed considerable loss of PGs almost throughout the tissue thickness when compared to both normal and moderate OA groups.

**Figure 2:**
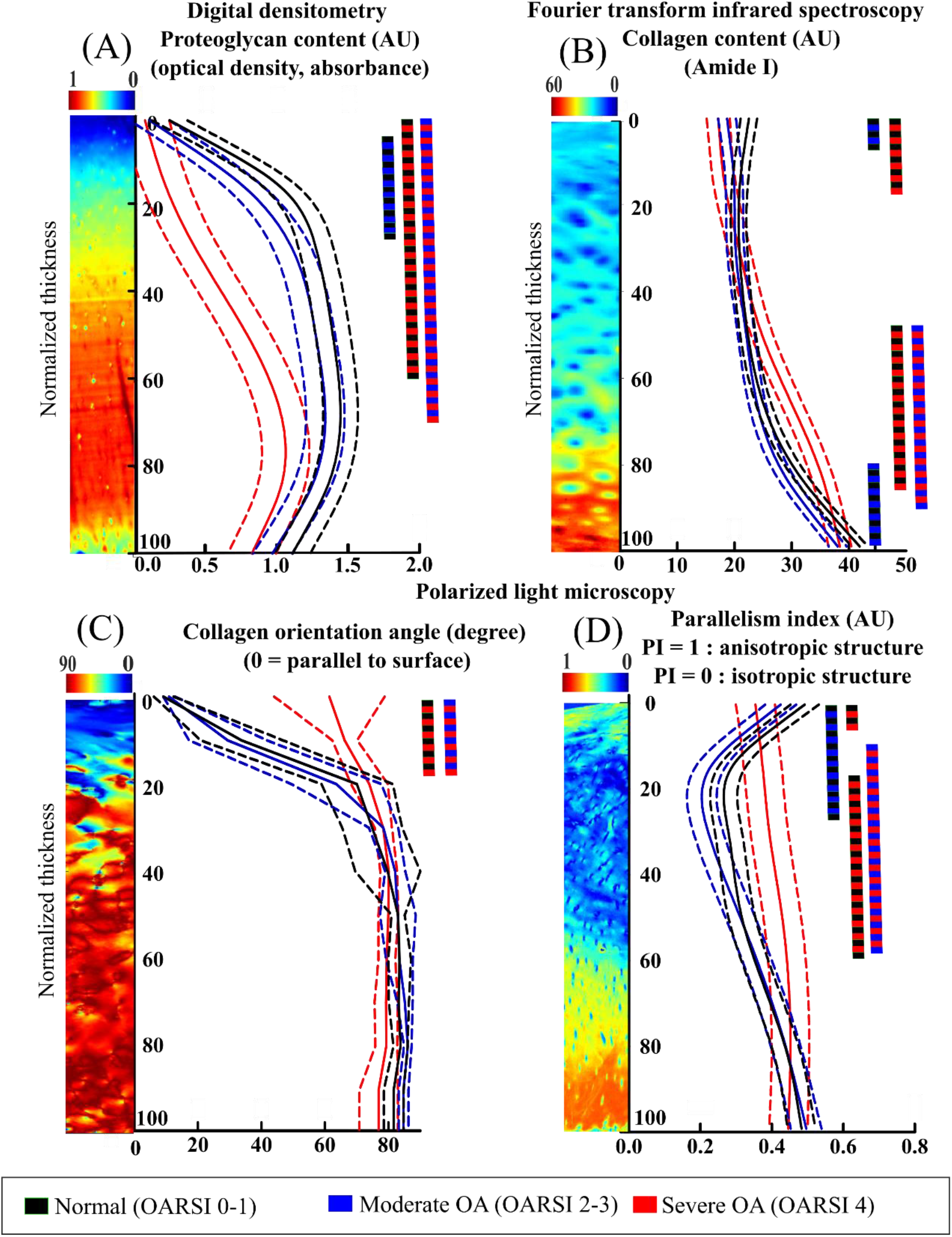
Depth-wise (A) PG content, (B) collagen content, (C) collagen orientation angle, and (D) anisotropy index in normal, moderate OA, and severe OA cartilage. The solid line shows the mean (median for the collagen orientation angle) and the dashed lines show ± 95% CI. Color bars indicate regions in which p < 0.05 (see Figure S1 for mean differences and 95% CI for the differences). The group-wise comparison of composition and structure in the deep cartilage near the tidemark is presented in Supplementary Figure S2.

The collagen content was smaller in the superficial layer (0-10% of tissue thickness) and in a region of the deep layer (77-100% of tissue thickness) of the moderate OA group compared to the normal group (Figure 2B). Surprisingly, the severe OA group had higher collagen content in the deepest layer compared to the normal and moderate OA groups.

The collagen orientation angle showed no essential difference between the normal and moderate OA groups (Figure 2C). However, the severe OA group had a substantially greater collagen orientation angle compared to the normal and moderate OA groups in the superficial layer (1-20% of tissue thickness).

The fibril parallelism index (anisotropy) was slightly smaller in the moderate OA group compared to the normal group (1-33% of tissue thickness, Figure 2D). However, the severe OA cartilage showed a significantly more uniform parallelism index profile (less variation as a function of the tissue depth) when compared to the normal samples (0-5% and 14-67% of tissue thickness) and moderate OA samples (10-64% of tissue thickness).

The mean differences and 95% CI for the differences for the structural and compositional parameters are presented in the supplementary material (Figure S1).

### 3.2. Depth-wise structure-function associations with the mechanical parameters

The structure-function relationships established based on the constituent-specific FRPE material parameters as well as the phase differences at 0.005 and 1 Hz loading frequency are shown in table 1 for the superficial (0-10%) and bulk tissue (0-100%). The PG content of the superficial layer showed a direct relationship with the initial fibril network modulus and non-fibrillar matrix modulus, while the PG content of the superficial layer was inversely associated with the strain-dependent fibril network modulus and the phase difference measured at 0.005 Hz. The PG content of the bulk tissue was directly associated with the non-fibrillar matrix modulus and the phase difference measured at 0.005 Hz (Table 1). The collagen orientation angle of the superficial layer showed an inverse association with the initial fibril network modulus while the collagen orientation angle of the bulk tissue was directly associated with the phase difference measured at 1 Hz. Interestingly, the permeability and its strain-dependency coefficient were not associated with any compositional and structural properties of the superficial layer or bulk tissue (Table 1). Further, the fibril parallelism index and collagen content were not associated with any functional property.

**Table 1:**
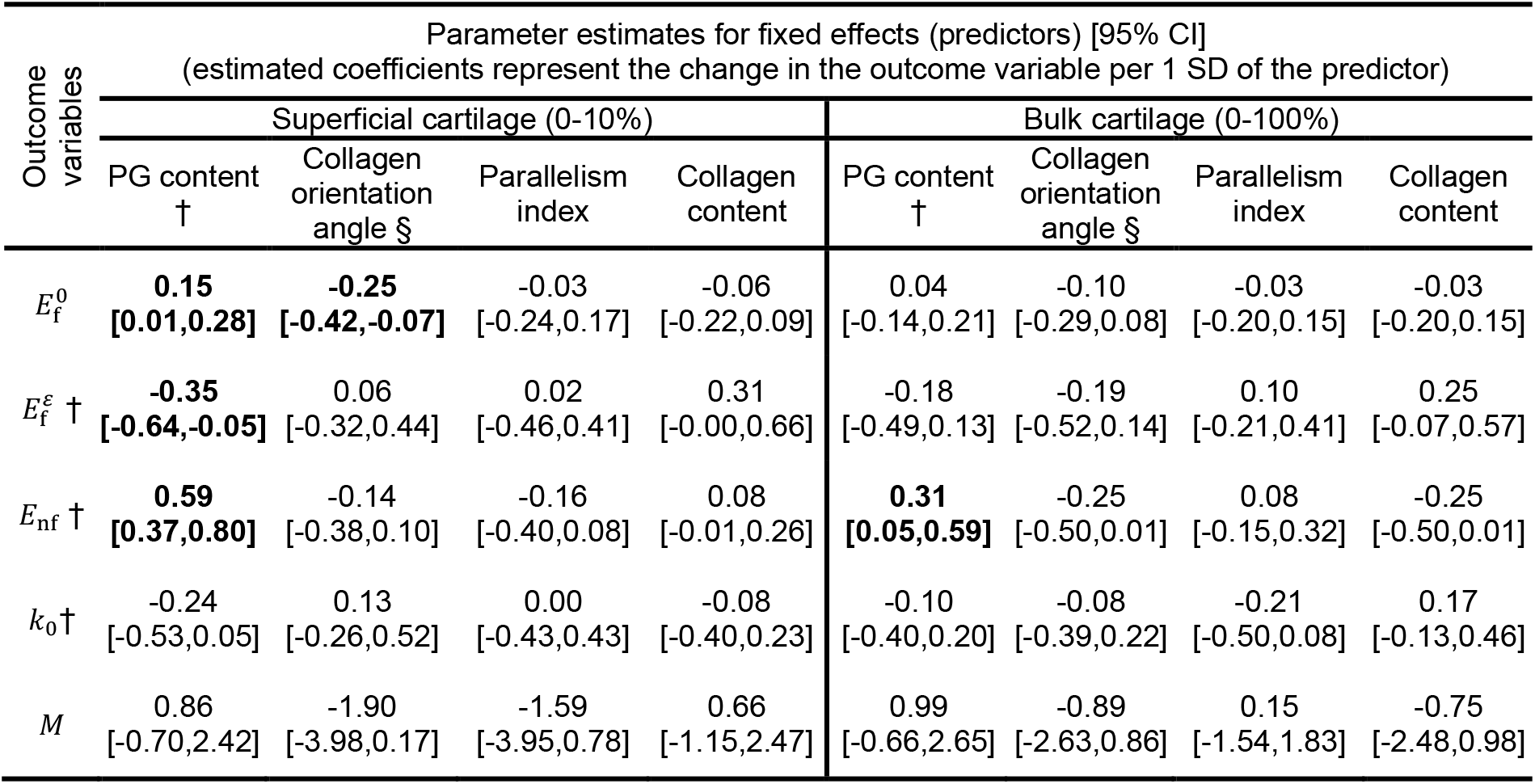
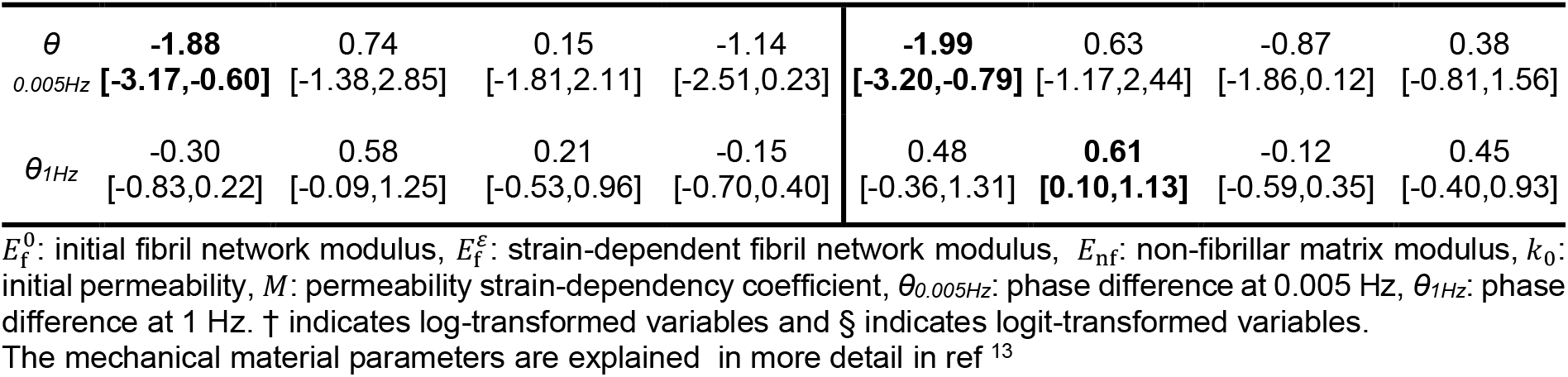
Structure-function relationships between the structural, compositional and FRPE material properties as well as viscosity (phase difference) in the superficial (0-10% of the normalized tissue thickness) and bulk cartilage using a linear mixed effect model. Bold font highlights cases in which the model was able to explain the relationship between structure and function based on the 95% CI of the predictors.

## 4. Discussion

This is the first study to determine structure-function relationships in human femoral cartilage at different stages of osteoarthritis. We conducted this by characterizing the depth-wise structural and compositional properties (PG content, collagen content, collagen fibril orientation angle, and the degree of fibril parallelism) and relating them to the model-derived constituent-specific and viscoelastic mechanical material properties. By doing this, we obtained new insights into how each cartilage constituent contributes to the mechanical response of human femoral cartilage at different stages of OA. This information is useful for translational research and can improve the representativeness of computational multiscale models of the knee joint.

### 4.1. Depth-wise structural and compositional alterations in human cartilage at different stages of OA

We observed a minor loss of the superficial PG content, minor loss of very superficial and very deep collagen content, minor loss of the superficial parallelism index, and no essential changes in the collagen orientation angle of moderate OA cartilage compared to those of normal cartilage. However, in severe OA cartilage, we observed a major loss of PG content, loss of superficial collagen content, greater collagen content in middle-deep tissue, and disorganization of collagen fibers in the uppermost cartilage layers compared to those of normal cartilage. Previous studies have reported that the superficial proteoglycan content in femoral condyle cartilage may not substantially alter at early OA ^2,12^.

We also observed that the collagen content in severe OA cartilage was greater in the deep tissue when compared to those of healthy and moderate OA cartilage. It should be noted that severe OA samples, especially OARSI 5 and 5.5 samples, had presumably lost their superficial zone, thus, the comparison between OA groups using the normalized thickness may not be fully reliable. Therefore, we also performed the group-wise comparison in the deep cartilage using absolute thickness values. In this analysis, the analysis of composition and structure started from the tidemark and extended 250 µm towards the cartilage surface. This analysis did not show essential differences in the composition or structure of deep cartilage between the groups (see supplementary material, Figure S2).

We want to emphasize that our observations hold only on the extracellular matrix level, and changes in composition (and function) may have already occurred in the cell microenvironment, as suggested by recent studies with human and mouse cartilage^6,32^. It must be also noted that our moderate OA group included mostly OARSI 2 (*n* = 16) than OARSI 3 grades (*n* = 3), thus the moderate OA group represents more mildly degenerated cartilage and the lack of differences in the composition and structure can be expected.

### 4.2. Associations between the mechanical material properties, structure and composition

Superficial PG degradation was associated with the smaller initial fibril network modulus. This was an interesting finding as in our previous study, which focused on biomechanical properties of the same samples, we did not observe changes in the initial fibril network modulus between normal and moderate OA cartilage (Figure 3). We also observed that all initial fibril network modulus values were substantially smaller in severe OA cartilage when compared to those of normal or moderate OA cartilage. One limitation in that study was that we had only 3 samples in the severe OA group, which made statistical evaluation impossible. Previously, we have suggested that the initial fibril network modulus represents the pretension of the collagen network. PG loss is also known to reduce the swelling pressure of cartilage ^51,52^. Thus, our result may also imply that the loss of PGs and tissue swelling pressure ultimately impairs the pretension of the collagen network in cartilage.

**Figure 3:**
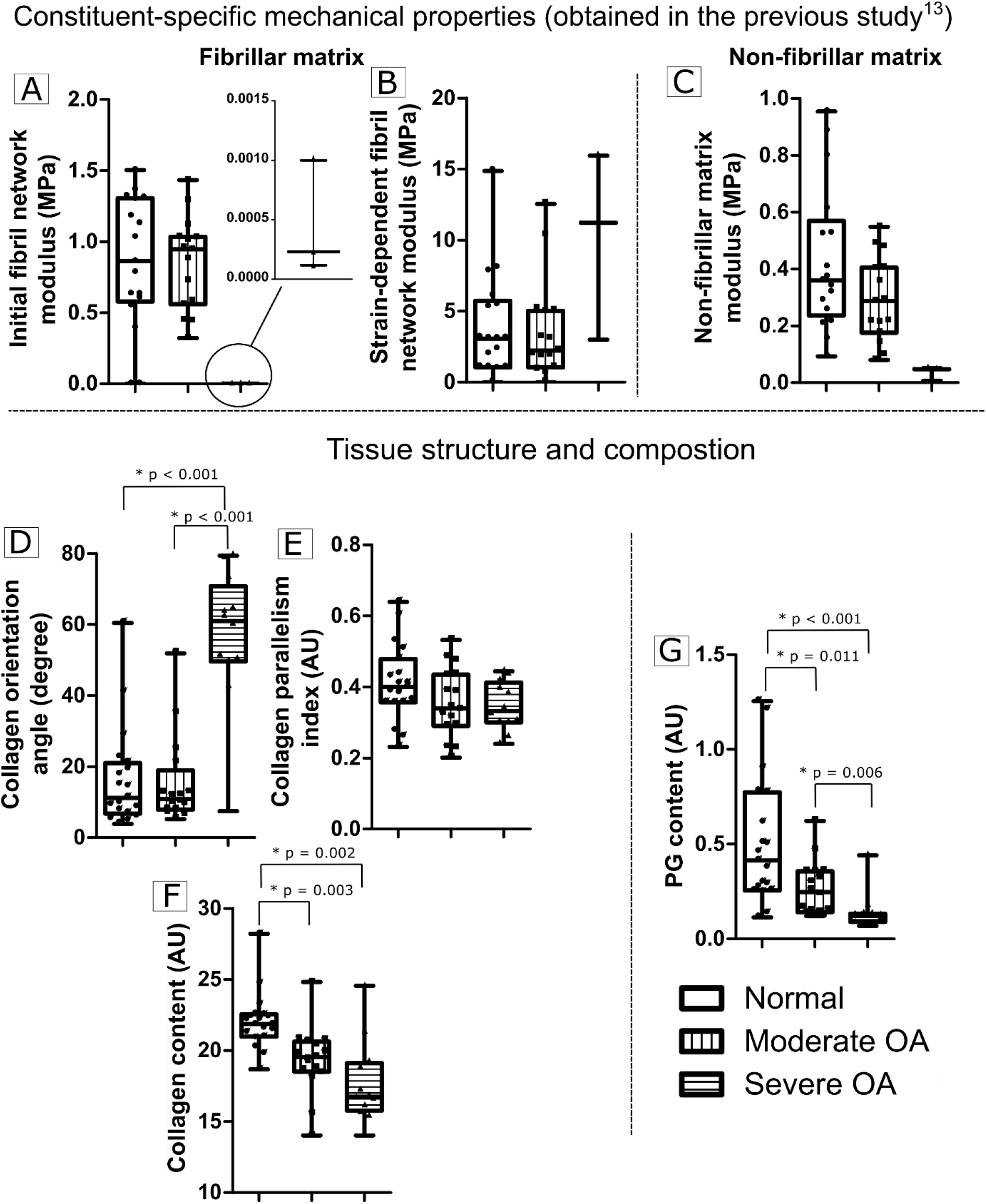
The summary of the observed changes in the function (A, B and C) and the structure (D, E, F and G) and of cartilage at different stages of OA. Boxplots show median, first quartile, third quartile and range of values. The results suggest that the loss of collagen pretension (= the initial fibril network modulus) at severe stages of OA is explained by the changes in the composition (PG content, G) and structure (collagen orientation, E) of the superficial cartilage. In severe OA cartilage, loss of collagen pretension (A) is regulated by both the PG content (G) and collagen disorganization (D). The results also suggest that the smaller non-fibrillar matrix modulus (C) of cartilage are explained by the loss of PG content (G) of the superficial cartilage in severe OA cartilage compared to normal cartilage.

Moreover, superficial PG degradation was associated with the greater nonlinear mechanical response of collagen fibrils (as indicated by the greater strain-dependent fibril network modulus). As with the initial fibril network modulus, the strain-dependent fibril network modulus did not change between normal and moderate OA cartilage in our previous biomechanical study (with the same samples, Figure 3). Further, severe OA cartilage showed signs of substantially greater strain-dependent fibril network modulus compared to those of normal and moderate OA cartilage, but we had the same issue with sample size as in the case of initial fibril network modulus. However, the relationship between strain-dependent fibril network modulus and PG content was not evident in human tibial cartilage, presumably due to dominant OA samples ^15^. In normal cartilage, swelling pressure is at healthy (or “normal”) levels and that separates, stretches, and prestresses the collagen fibrils (i.e. they are less crimped) ^1,33^. This results in a less nonlinear mechanical response of the collagen network. However, in the more degenerated tissue, the swelling pressure is reduced, thus, the fibrils in severe OA cartilage are less prestressed and more crimped. Thus, at the initial phase of loading, fibrils need to be straightened first before their mechanical contribution is fully employed. This phenomenon is seen in our results as a low initial fibril network modulus and a high strain-dependent fibril network modulus in severe OA cartilage.

As expected, superficial (and bulk) PG degradation was associated with the smaller non-fibrillar matrix modulus. We also observed a similar kind of trend in OA group-wise comparisons (Figure 3). Similar associations have been shown in several animal and human cartilage studies ^15,25^, and while this is not a new result, it helps in verifying the used methods.

Interestingly, the disorganization of the collagen network (greater collagen orientation angle) in superficial cartilage was also associated with the smaller initial fibril network modulus. Based on group-wise comparisons, the disorganization was evident only in severe OA cartilage and specifically in superficial tissue (Figure 2c and 3). This relationship was evident in human tibial cartilage as well ^15^. Previous in vitro studies (and our current results) suggest that the PG loss may precede the changes that occur in the collagen network ^23,26^, thus this association between collagen disorganization and smaller initial fibril network modulus may be explained by the superficial PG loss. This was also reflected in seemingly unchanged fibrillar matrix parameters of moderate OA and normal cartilage samples. It is also possible that the collagen fibril network in severe OA cartilage experiences alterations in its molecular level structure (e.g., fibrils or crosslinks are damaged and/or cleaved due to proteolytic enzyme activity ^20^) which may impair the capability of the collagen fibrils to bear pre-tension.

The loss of PGs was associated with a more viscoelastic (i.e., greater phase difference) mechanical response at the low loading frequency (0.005 Hz). This was in contrast with our previous observations in tibial cartilage. In that study, we observed that collagen content, instead of the PG content, was associated with the phase difference at 0.005 Hz. Thus, in our previous study, we speculated that the inherent viscoelasticity of collagen might be one of the main contributors to the apparent viscoelastic response. Yet, in light of our current results, we deduct the following explanation from our observations. In normal cartilage, the PG content is high and the fluid flow is more restricted due to small permeability. Thus, the mechanical behavior of normal cartilage is more elastic compared to the more osteoarthritic cartilage, in which the lack of PGs may facilitate fluid flow and that is seen as greater apparent tissue viscoelasticity (phase difference). It is also known that cartilage viscosity is sensitive to biochemical alterations occurring in the matrix constituents as, for instance, proteoglycan depletion causes a reduction in the energy storage capacity of the matrix ^21,37^. Interestingly, the collagen disorganization, rather than PG loss, was associated with a more viscoelastic mechanical behavior at the high loading frequency (1 Hz). This could indicate that the disorganized collagen network of more osteoarthritic samples is not as capable of trapping and pressurizing the interstitial fluid compared to the normal tissue with a properly organized collagen network. This is also supported by an earlier computational study suggesting that the proper arrangement of the collagen network plays an important role in restricting and directing fluid flow ^17,41^.

It is known that the apparent viscoelastic response of cartilage is a combination of fluid flow-dependent (fluid-solid interactions, permeability i.e., poroelasticity) and flow-independent (inherent viscoelasticity of PGs and collagen) mechanisms ^24,30,41^. To further elucidate what is the contribution of the flow-dependent mechanism, and how that alters in OA, in our experiments, we simulated FE models with median FRPE material parameter values of normal and severe OA cartilage and extracted phase differences and fluid pressure at different loading frequencies (Figure 4). We also checked that our analysis remains valid when we select sample-specific material parameters for representative normal and severe OA samples (see Figure S3). According to these simulations, the poroelastic contribution (i.e., phase difference peak) is greatest at around 0.0005 and 0.005 Hz for normal and severe OA cartilage. This suggests that the apparent viscoelastic material response originates mainly from the flow-dependent mechanism (poroelasticity) in our experiments and the maximum contribution of poroelasticity is not even reached in normal cartilage. Further, the FRPE model predicted phase difference at 1 Hz was ∼0.7 and 2.9 degrees in normal and severe OA cartilage, suggesting that fluid flow has a very small contribution to the apparent viscoelastic response at these frequencies (i.e., fluid is trapped inside cartilage, and it is not able to flow which results in fluid pressurization as seen in Figure 4b). Interestingly, our experimental data and other studies suggest that healthy and OA cartilage has substantially greater phase difference at 1 Hz loading frequency than what the model predicted (median difference between experiment and model ∼8 and ∼10 degrees for normal and severe OA cartilage). The fluid flow-dependent viscoelasticity is length scale dependent ^19,36,37^ while flow-independent apparent viscoelasticity is not (i.e., phase difference peak remains at the same frequency)^39^. Earlier atomic force microscopy-based nanorheology studies have also suggested that flow-independent mechanism(s) may have a major contribution to the phase difference of cartilage at the frequency range of ∼1-10 Hz ^36,39,50^. As the FRPE model did not include any flow-independent mechanisms, the differences between the model-predicted and experimentally observed phase differences should originate from the flow-independent viscoelasticity of collagen and/or collagen-PG interactions. They can also partially originate from measurement uncertainties. We aim to investigate this crosstalk between flow-dependent and independent mechanisms and how they alter at different stages of OA in more detail in a future computational study.

**Figure 4:**
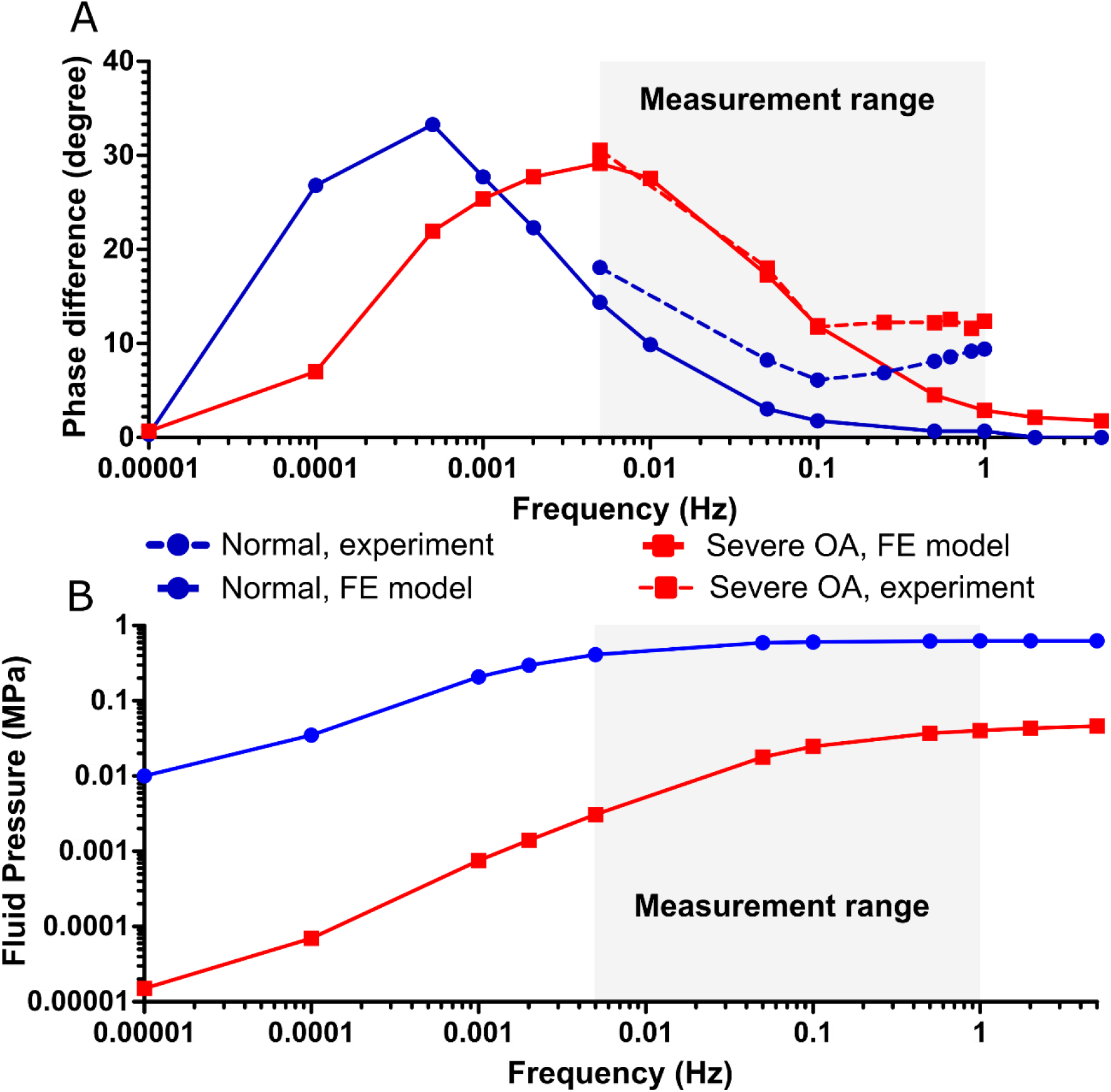
The phase difference (A) and fluid pressure (B) extracted from FE models with median FRPE material parameter values of normal and severe OA cartilage. The median FRPE material parameters for the normal sample 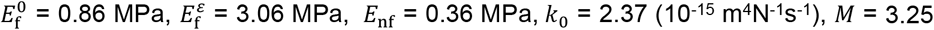 and for the severe OA sample 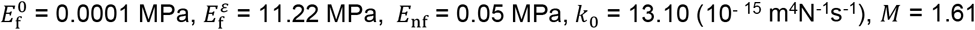.

The tissue permeability was neither substantially different between normal and OA groups nor it could be explained by the structural or compositional alterations at different stages of OA. This was surprising as previous studies have suggested that both the collagen fibril network and glycosaminoglycan side chains of PGs modulate the permeability of cartilage ^17,28,43^. Possible explanations for this can be 1) our sample set was dominated by normal and moderate OA samples and 2) the moderate OA samples seem to have maintained (as suggested by our results) the normal condition of their collagen network. Thus, the tissue permeability was not altered enough in our dataset which is why we could not relate it to the structure and composition of cartilage.

### 4.3. Limitations

We pooled the OARSI grades 0-1.5, OARSI grades 2-3.5 as well as OARSI grades 4-5.5 into the normal, moderate OA and severe OA groups, respectively, to obtain a meaningful number of samples in each group. The groups were not balanced in their OARSI score distribution, as the moderate OA group tends to represent OARSI 2 (i.e., few OARSI 3 samples), which could lead to smaller differences than if the moderate group would have contained more OARSI grade 3 samples.

To obtain the depth-dependent properties of the tissue, structural and compositional properties were normalized to the tissue thickness. However, it is known that the superficial layer is often lost at the very late stages of OA ^16^. On the other hand, tissue thickness has also been observed to increase due to the disorganization of the collagen network and cartilage swelling ^4,9^ in early OA. These differences in cartilage thicknesses could potentially affect our depth-wise analyses, especially with very severe OA samples, e.g. OARSI 5-6 ^42^. Our femoral condyle samples included samples scored as OARSI 5, thus the point-by-point comparison of depth-wise compositional properties of this group to the other groups may not be fully reliable. Moreover, we cannot say for certain whether the tissue has lost any layers or not (as we did not follow the disease progression of the samples). However, the established structure-function relationships of human cartilage should be plausible because 1) no samples with considerable tissue loss (i.e., OARSI 5) were included in structure-function analyses, 2) biomechanical testing and analysis of tissue composition and structure were consistently performed to the normalized tissue thickness. In addition, our severe OA group consisted of only medial samples, while the normal and moderate OA groups consisted of both medial and lateral samples. Although we did adjust the analyses for knee compartment, we could not assess compartment-specific between-group differences for the severe OA group.

### 4.4. Conclusion

In conclusion, we presented novel information on structure-function relationships in human femoral condyle cartilage. We observed that 1) the PG degradation (which is known to lead to loss of tissue swelling pressure) and 2) disorganization of the collagen fibril network in superficial cartilage reduced the pretension of the collagen fibrils. Moreover, the PG degradation in superficial cartilage increased the nonlinear mechanical response of the collagen fibrils. PG degradation also modulated tissue viscosity at the low loading frequency, while the disorganization of the collagen network modulated tissue viscosity at a high loading frequency.

## Conflicts of interest

M Englund declares serving as panel member on an advisory board for Pfizer (tanezumab, Nov 2019).

## Acknowledgments

This work was supported by Academy of Finland (grants 286526, 324529 and 268378); strategic funding of the University of Eastern Finland; Maire Lisko Foundation; Finnish Cultural Foundation, North Savo Regional Fund (grant 65191841); Emil Aaltonen Foundation (grant 200016); and Alfred Kordelin Foundation (grant 190111). The study (MENIX biobank) and contributions by M Englund and A Turkiewicz is supported by the Swedish Research Council, The Swedish Rheumatology Association, Österlund Foundation and Governmental funding of clinical research within the national health services (ALF).

## Notes

### Competing Interest Statement

Martin Englund declares serving as panel member on an advisory board for Pfizer (tanezumab, Nov 2019).

