## Supplementary Table S1 for "Associations of human femoral condyle cartilage structure and composition with viscoelastic and constituent-specific material properties at different stages of osteoarthritis"

**Results**

Table S1: Descriptive statistics on samples used to estimate differences in structural parameters between OA-groups (n = 47).

|  | OARSI grade | Medial donor | Medial TKR | Lateral donor | Lateral TKR | Age  (mean ± SD) | BMI  (mean ± SD) | Sex  (M or F) |
| --- | --- | --- | --- | --- | --- | --- | --- | --- |
| Healthy | 0 | 1 | - | 2 | 2 | 49.3 ± 19.5 | 28.5 ± 7.6 | 2M, 3F |
|  | 1 | - | - | - | 1 | 66 | 35 | 1F |
|  | 1.5 | 4 | - | 2 | 7 | 57.2 ± 15.1 | 28.8 ± 6.6 | 7M, 6F |
| Moderate OA | 2 | 2 | - | - | - | 59.5 ± 1.5 | 28.0 ± 5.0 | 1M, 1F |
|  | 2.5 | 3 | - | 3 | 5 | 65.0 ± 14.1 | 26.8 ± 3.5 | 5M, 6F |
|  | 3 | - | - | 1 | - | 70 | 25 | 1M |
|  | 3.5 | - | - | 2 | - | 59.5 ± 1.5 | 28.0 ± 5.0 | 1M, 1F |
| Severe OA | 4.5 | - | 5 | - | - | 65.0 ± 7.6 | 29.6 ± 5.3 | 1M, 4F |
|  | 5 | - | 6 | - | - | 63.0 ± 7.4 | 32.5 ± 3.3 | 2M, 4F |
|  | 5.5 | - | 1 | - | - | 67 | 26 | 1M |

Table S2: Descriptive statistics on the subset of the samples used to estimate associations between structural and biomechanical parameters (n = 35, note that out of 47 samples, 12 were not measurable in the mechanical indentation test).

| OARSI grade | Medial donor | Medial TKR | Lateral donor | Lateral TKR | Age  (mean ± SD) | BMI  (mean ± SD) | Sex  (M or F) |
| --- | --- | --- | --- | --- | --- | --- | --- |
| 0 | 1 | - | 2 | 2 | 49.3 ± 19.5 | 28.5 ± 7.6 | 2M, 3F |
| 1.5 | 4 | - | 2 | 6 | 55.8 ± 15.1 | 28.7 ± 6.9 | 6M, 6F |
| 2 | 2 | - | - | - | 59.5 ± 1.5 | 28.0 ± 5.0 | 1M, 1F |
| 2.5 | 3 | - | 2 | 5 | 66.6 ± 14.1 | 26.8 ± 3.7 | 4M, 6F |
| 3 | - | - | 1 | - | 70 | 25 | 1M |
| 3.5 | - | - | 2 | - | 59.5 ± 1.5 | 28.0 ± 5.0 | 1M, 1F |
| 4.5 | - | 3 | - | - | 70.3 ± 3.7 | 33.3 ± 3.1 | 1M, 2F |

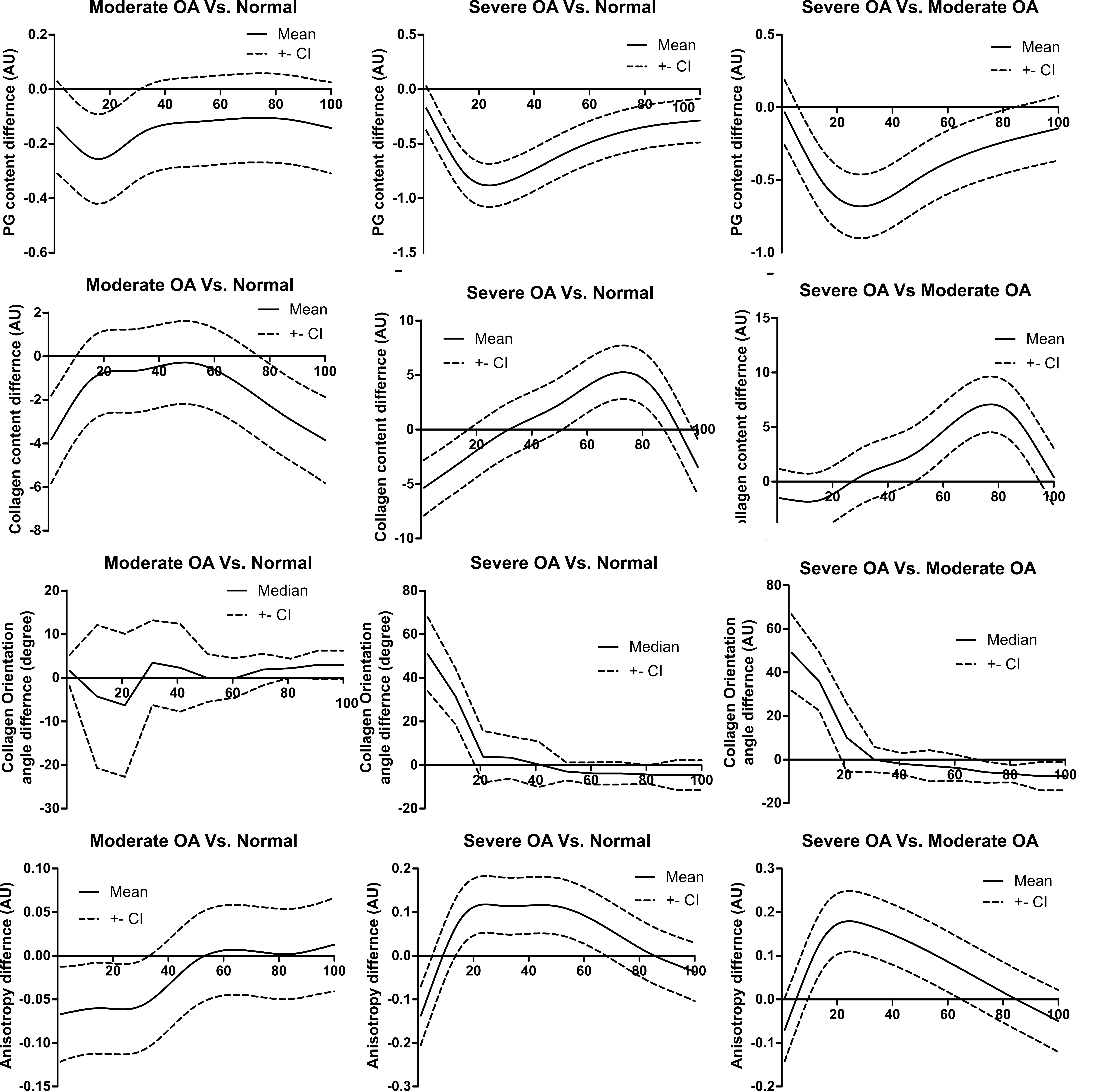

Figure S1: Differences for the depth-wise PG content, collagen content, collagen orientation angle and collagen fibril parallelism index (anisotropy) between normal, moderate OA, and severe OA groups. The solid line represents the mean (or median for collagen orientation) difference and the dotted line represents the 95% confidence intervals (CI).

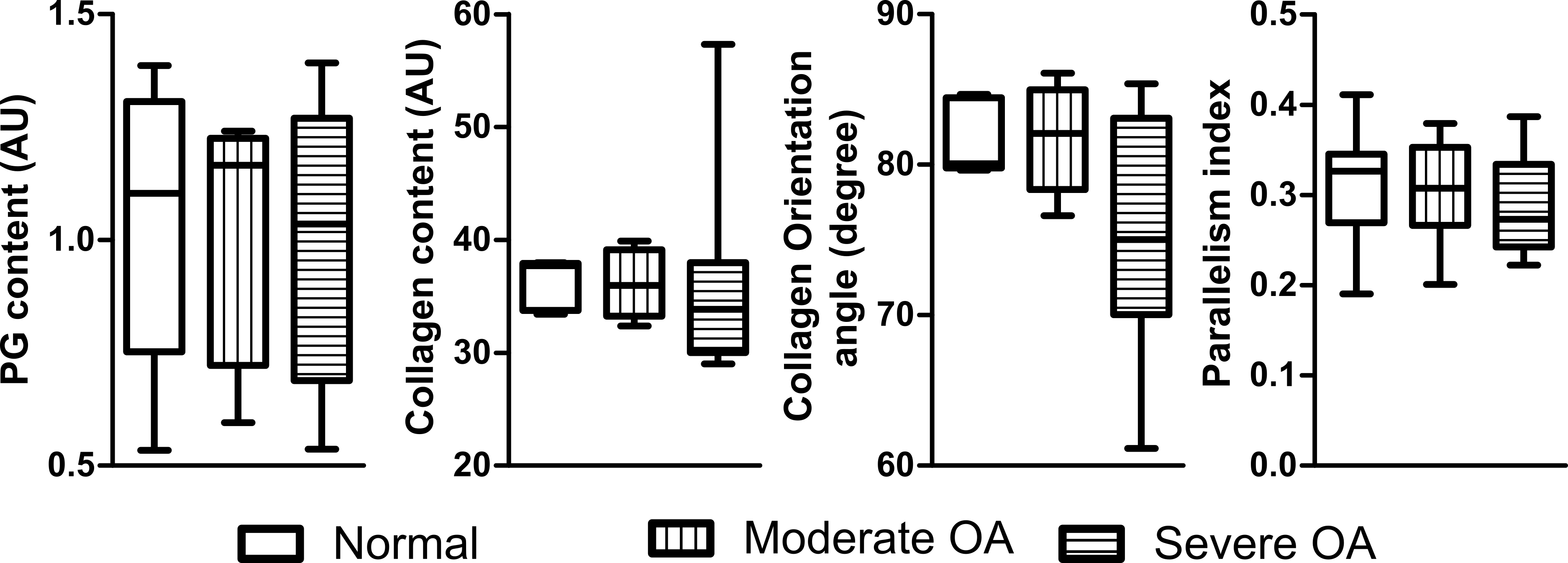

Figure S2: PG content, collagen content, and collagen orientation angle of the deep cartilage in normal, moderate OA, and severe OA groups. The analysis was conducted by averaging a 250 µm thick region starting from the tidemark towards the surface.

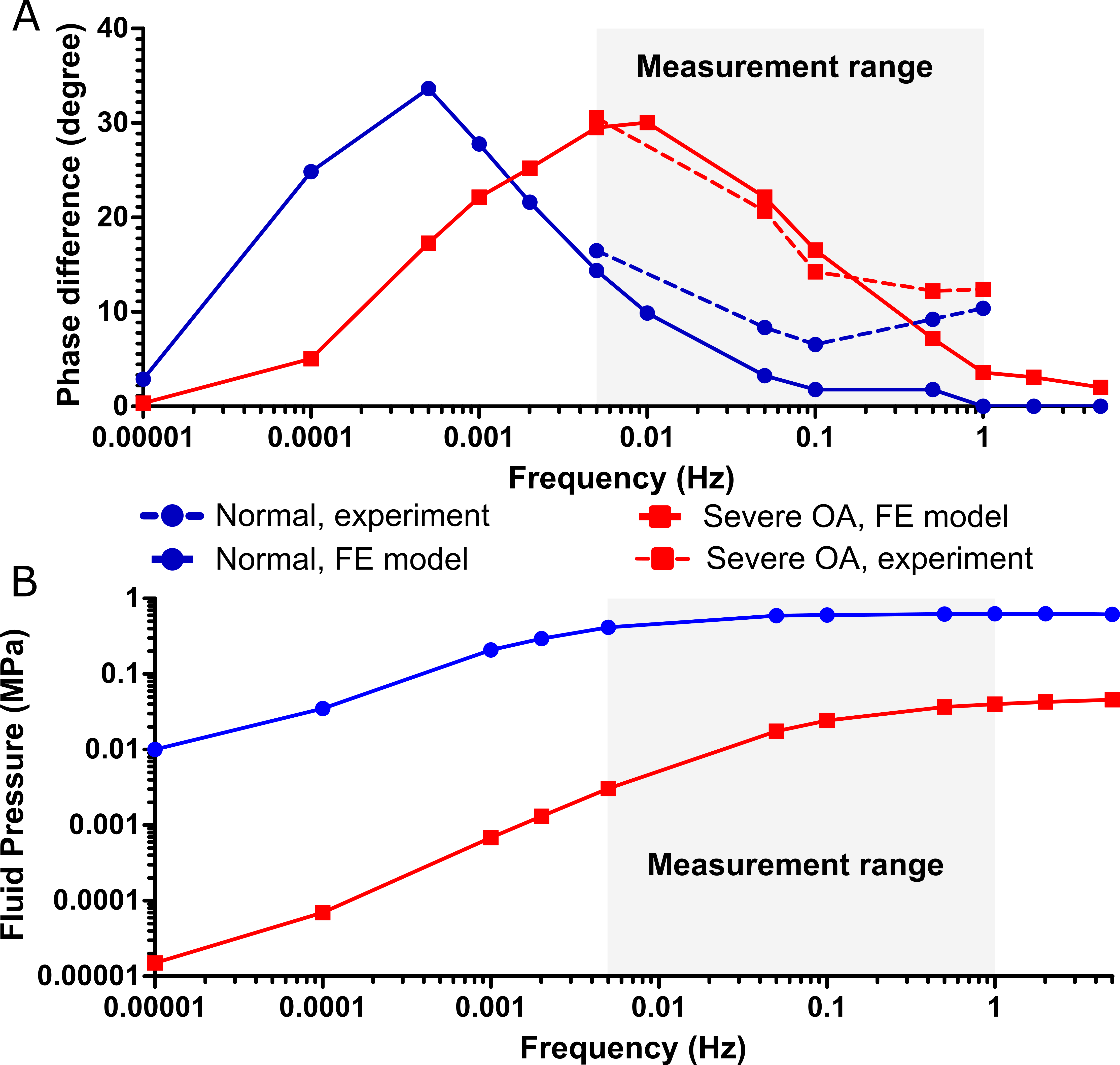

Figure S3: Experimental and FRPE model predicted sample-specific A) phase difference and B) fluid pressure as a function of frequency for a representative normal and severe OA human femoral cartilage. The experimental measurement range is shown as a shaded region. The FRPE material parameters for the normal sample $E_{f}^{0}$ = 1.30 MPa, $E_{f}^{\varepsilon}$ = 1.04 MPa, $E_{\mathrm{nf}}$ = 0.53 MPa,$k_{0}$ = 2.04 (10^-15^ m^4^N^-1^s^-1^), $M$ = 8.88 and for the severe OA sample $E_{f}^{0}$ = 0.0002 MPa, $E_{f}^{\varepsilon}$ = 11.22 MPa, $E_{\mathrm{nf}}$ = 0.05 MPa,$k_{0}$ = 13.10 (10^-15^ m^4^N^-1^s^-1^), $M$ = 1.61.
